# Protein nonadditive expression and solubility in *Arabidopsis* hybrids and allotetraploids

**DOI:** 10.1101/2023.03.01.530688

**Authors:** Viviana June, Dongqing Xu, Ophelia Papoulas, Daniel Boutz, Edward M. Marcotte, Z. Jeffrey Chen

## Abstract

Interspecific hybridization in plants often leads to allopolyploids including most important crops such as wheat, cotton, and canola, while many other crops such as corn and sorghum are grown as hybrid. Both allopolyploids and hybrids show hybrid vigor or heterosis. The phenomenon has been widely applied in agriculture and extensively studied at the genetic and gene expression levels. However, proteomic changes in hybrids and polyploids remain poorly understood. Here, we report comparative analysis of soluble and insoluble proteomes in *Arabidopsis* intraspecific and interspecific hybrids or allotetraploids formed between *A. thaliana* and *A. arenosa*. Both allotetraploids and intraspecific hybrids displayed nonadditive expression (unequal to the sum of the two parents) of the proteins, most of which were involved in biotic and abiotic stress responses. In the allotetraploids, homoeolog-expression bias was not observed among all proteins examined but could occur among 17-20% of the nonadditively expressed proteins, consistent with the transcriptome results. Among expression-biased homoeologs, there were more *A. thaliana*-biased than *A. arenosa*-biased homoeologs,. Analysis of the insoluble and soluble proteomes revealed more soluble proteins in the hybrids than their parents but not in the allotetraploids. Most proteins in ribosomal biosynthesis and in the thylakoid lumen, membrane, and stroma were in the soluble fractions, indicating a role of protein stability in photosynthetic activities for promoting growth. These results collectively support roles of nonadditive expression of stress-responsive proteins and increased solubility of photosynthetic proteins in *Arabidopsis* hybrids and allotetraploids, which may promote hybrid vigor.

**Plain Language Summary:** We analyzed fractionated proteomes to test protein abundance and solubility in *Arabidopsis* hybrids and polyploids. Many proteins in stress-responses are nonadditively expressed in intraspecific hybrids or allotetraploids. There are more soluble proteins of ribosomal biosynthesis and photosynthetic activities in the hybrids than in their parents but not in allotetraploids. Expression levels are equal among most protein homoeologs in the allotetraploids but are biased for ~20% of nonadditively expressed proteins. Nonadditive protein expression and solubility may play a role in heterosis.

## Introduction

Heterosis or hybrid vigor refers to the observation that hybrid offspring show greater growth and fitness than either parent and occurs across plant and animal kingdoms. phenomenon was systematically described by Charles Darwin in 1876 (Darwin 1876), and rediscovered by Shull and East during maize breeding (Darwin 1876; East 1936; Shull 1908). Several genetic models are available to explain heterosis. The dominance model suggests complementation of deleterious alleles by the dominant ones in the heterozygous loci (Bruce 1910; Jones 1917). The overdominance model indicates that heterozygous loci in hybrids are expressed at a higher level than or advantageous over homozygous loci (Crow 1948; East 1936). Another model is related to epistasis, in which interactions between nonallelic genes contribute to the growth vigor in hybrids (Schnell and Cockerham 1992; Yu *et al*. 1997). However, no single model can fully explain the basis of heterosis.

A notion in the field is to jump outside theoretical dogmas because these genetic models cannot address epistasis or complex regulatory network interactions in various biological pathways (Birchler *et al*. 2010). Indeed, transcriptomic analyses have revealed genome-wide nonadditive gene expression changes in *Arabidopsis* allotetraploids or interspecific hybrids (Wang *et al*. 2006b), which led to the discovery of linking enhanced circadian rhythms with biomass heterosis in plant hybrids (Ni *et al*. 2009). Expression peaks of circadian clock genes are epigenetically altered in the hybrids to enhance expression of the circadian output genes in photosynthesis and starch biosynthesis. The more starch is synthesized during the day, the more it can be degraded at night to promote growth (Chen 2013). The role of altered circadian rhythms in heterosis has been consistently demonstrated in *Arabidopsis* (Shen *et al*. 2012), rice (Chen *et al*. 2010), and maize (Ko *et al*. 2016; Li *et al*. 2020), suggesting a conserved role of enhanced circadian rhythms in hybrid vigor.

Studies of proteomic changes in hybrids are very limited. Using protein two-dimensional gel electrophoresis analysis of the proteins extracted from mitochondria, Dahal *et al*. found a correlation between expression of specific alleles and/or post-translational modification of specific proteins and higher levels of heterosis in different maize hybrids (Dahal *et al*. 2012). Using isobaric tags for relative and absolute quantitation (iTRAQ) coupled with mass spectrometry, Ng *et al*. found that expression of ~8% of the proteins in *Arabidopsis* allotetraploids are nonadditive relative to the parents (mid-parent level) (Ng *et al*. 2012). Although the overall trend of nonadditive expression is consistent between transcript and protein levels, the percentage of differentially accumulated proteins that matched differentially expressed genes is relatively low. In natural allopolyploid *Tragopogon mirus*, hybridization generates more effects on proteomes than polyploidy (Koh *et al*. 2012). In maize hybrids, metabolic changes correspond to nonadditive protein abundance and enzyme activities of key enzymes in the respective pathways, suggesting that concerted changes in metabolomes and proteomes contribute to maize heterosis (Li *et al*. 2020). Another study indicates increased expression of nuclear- and plastid-encoded subunits of protein complexes required for protein synthesis in chloroplasts and for photosynthetic activities in hybrid seedling leaves, and hybrid/mid-parent expression ratios of chloroplast ribosomal proteins are correlated with plant height heterosis (Birdseye *et al*. 2021). These results suggest that post-transcriptional regulation and protein synthesis play a role in regulating the nonadditive expression of proteins in hybrids (Ng *et al*. 2012; Yang *et al*. 2021).

Metabolic and proteomic studies in maize further demonstrate that a large fraction of maize metabolites and proteins is diurnally regulated, and many show nonadditive abundance in the hybrids (Li *et al*. 2020). Metabolic heterosis is relatively mild, and metabolites in the photosynthetic pathway show positive mid-parent heterosis (MPH), whereas metabolites in the photorespiratory pathway show negative MPH. Hybrids may more effectively remove toxic metabolites generated during photorespiration, and thus maintain higher photosynthetic efficiency for heterosis. The cause of these changes remains elusive. One possibility is that the presence of multiple different alleles of a single gene in hybrids allows for selective expression of the more stable alleles (Goff 2011).

Here, we investigated both changes in protein abundance and solubility in two sets of hybrids: reciprocal intraspecific hybrids between *Arabidopsis thaliana* ecotypes C24 and Col-0 (Miller *et al*. 2015), and allotetraploids *A. suecica* and Allo738 and their progenitors *A. arenosa* and *A. thaliana* (Jiang *et al*. 2021; Wang *et al*. 2006b). *A. thaliana* intraspecific hybrids have been extensively used as a model to study heterosis, as several hybrids (including Col/C24 hybrids) display high levels of growth vigor (Chen 2013; Groszmann *et al*. 2014). However, the parental ecotypes have similar genomes with fewer non-synonymous mutations compared to interspecific hybrids or allotetraploids, which also display increased levels of heterosis (Chen 2010; Chen 2013). A comparison of proteome changes between allotetraploids and intraspecific hybrids would allow for testing the effect of genetic distance on protein changes.

We applied a protein fractionation approach coupled with label-free liquid chromatography-mass spectrometry (LC-MS) to investigate proteomic changes in the intraspecific hybrids and allotetraploids. We found nonadditive expression of proteins in stress response, photosynthesis, and protein biosynthesis in the hybrids and allotetraploids, which are consistent with transcriptome results related to heterosis. There were more soluble proteins in the intraspecific hybrids relative to the parents, but not in the allotetraploids. Most ribosomal proteins and proteins in the thylakoid lumen, membrane, and stroma, were in the soluble fractions. These results may suggest a role of nonadditive regulation of stress-responsive and photosynthetic proteins in heterosis. Alternatively, reduced levels of protein synthesis may contribute to growth vigor in the hybrids and allotetraploids.

## Methods

### Plant materials

Two *A. thaliana* ecotypes Columbia (Col-0) and C24 were used as parents to generate reciprocal intraspecific hybrids by manually crossing as previously described (Miller *et al*. 2012). Each parent was also manually crossed as a control. Seeds were collected from these crosses once siliques had matured. Allotetraploid Allo738 was derived from an induced autotetraploid *A. thaliana* L*er* ecotype (Ath4; ABRC CS3900) and *A. arenosa* (Aar, Care-1; ABRC; CS3901), an outcrossing tetraploid species (Comai *et al*. 2000). Natural allotetraploid *A. suecica* strain As9502 (As; ABRC CS22509) and all other parental strains (Ath4 and Aar) were maintained in the lab.

### Plant growth conditions

Seeds were sterilized in 20% bleach for 10 minutes, followed by five rinses with 1 mL sterile ddH_2_O. Seeds were then plated onto 0.5 Murashige and Skoog media supplemented with 1% sucrose and stratified at 4°C in the dark for 48 hours. After stratification, seeds were transferred to a 22°C growth room with 16 hours of light and 8 hours of dark per day. Seven days after germination, seedlings were transplanted onto soil. A 3:1 mixture of Pro-Mix Biofungicide to Field and Fairway was used, and at first watering, plants were treated with 4g Miracle Gro Plant Food and 1 tsp Gnatrol Biological Larvicide (Valent Biosciences LLC, Libertyville, IL) per gallon of water. Plants were sprayed with Bonide copper soap fungicide weekly to prevent powdery mildew infection and with pesticide weekly to prevent thrips infestation.

### Protein extraction and fractionation

At 21 days after sowing, rosettes were harvested at zeitgeber time (ZT) 0 (dawn) to minimize circadian effects with 3 biological replicates for each genotype and flash-frozen in liquid nitrogen. A pool of 10 rosettes was ground to a fine powder in a chilled mortar and pestle. An equivalent volume of lysis buffer (50 mM Tris pH 7.5, 150 mM NaCl, 5 mM EGTA, 10% Glycerol, 1% NP40) with plant protease inhibitor cocktail (Sigma-Aldrich, St. Louis, MO) and phosphatase inhibitor (PhosSTOP Easy, Roche, Basel, Switzerland) was added to each sample. Samples were then lysed at 4°C on a rotator for 30 minutes. Debris was pelleted via centrifugation at 1,000 g for 10 minutes. The supernatant was retained as the whole cell extract. The whole cell extract was then fractionated into the soluble and insoluble fractions through centrifugation at 10,000 g for 10 minutes. The supernatant was retained as the soluble fraction, and the pellet was resuspended in lysis buffer to form the insoluble fraction. Fractions were then denatured in 50% trifluoroethanol (TFE) and 5 mM tris (2-carboxyethyl phosphine) (TCEP) at 55°C for 45 minutes. Samples were cooled to room temperature and alkylated in 15 mM iodoacetamide (IAM) at room temperature in the dark for 30 minutes. After the alkylation reaction was quenched with 7 mM dithiothreitol, the samples were diluted in trypsin digestion buffer (50 mM Tris, 2mM CaCl_2_, pH 8.0) to reduce the final TFE concentration to 5%. After adding 2 µg MS grade trypsin in the intraspecific hybrids (Pierce Biotechnology, Waltham, MA) and polyploids (Promega Corporation, Madison, WI) to each sample, the samples were digested at 37°C for 5 hours. Formic acid was added to a final concentration of 1% to quench the digestion. Sample volumes were reduced in a SpeedVac to 250 µL. Samples were then filtered using Amicon Ultra 10kD (Millipore Sigma, Burlington, MA) spin-caps to remove undigested protein and eluted in buffer C [95% H_2_O, 5% acetonitrile (ACN), 0.1% formic acid]. Samples were desalted using a 5-7 µL C18 Filter Plate (Glygen Corp.) and a vacuum manifold, eluted in 60% ACN, and reduced in volume to <10 µL in a SpeedVac. The final samples were resuspended in buffer C for mass spectrometry.

### Mass spectrometry

Mass spectra from each of three biological replicates were acquired on a Thermo Orbitrap Fusion Lumos. Peptides were separated using reverse phase chromatography on a Dionex Ultimate 3000 RSLCnano UHPLC system (Thermo Fisher Scientific, Waltham, MA) with a C18 trap to Acclaim C18 PepMap RSLC column (Dionex; Thermo Fisher Scientific) configuration. Peptides were eluted using a 5-40% acetonitrile gradient in 0.1% formic acid over 120 min for all samples. Peptides were injected directly into the mass spectrometer using nano-electrospray for data-dependent tandem mass spectrometry. The data acquisition used for the mass spectrometer was as follows: full precursor ion scans (MS1) collected at 120,000 m/z resolution. Monoisotopic precursor selection and charge-state screening were enabled using Advanced Peak Determination (APD), with ions of charge > +1 selected for high energy collision dissociation (HCD) with collision energy 30% stepped ± 3%. Dynamic exclusion was active with 20-second exclusion for ions selected twice within a 20 s window for intraspecific hybrid samples, and with 60 s exclusion for ions selected twice within a 60-second window for polyploid samples. All MS2 scans were centroid and done in rapid mode.

### Peptide assignment

For *A. thaliana*, the Arabidopsis proteome was downloaded from Uniprot in July 2018 (UniProt 2021). For the allotetraploids, the Allo738 proteome was generated from the recent long read resequencing of the Allo738 genome (Jiang *et al*. 2021). We then created an orthogroup collapsed proteome by concatenating the sequences of all proteins within orthogroups with triple lysines between each protein, as described in a published paper (McWhite *et al*. 2020). Orthogroups used to create the proteome were those identified in a previously published paper (Jiang *et al*. 2021). Peptide assignment was performed using Proteome Discoverer (v2.3 for the allotetraploid, and v2.2 for the *A. thaliana* hybrids). The MS spectra were searched against these proteomes as well as a database of common contaminants from MaxQuant using the SEQUEST HT node. For the search, a maximum of two missed trypsin cleavage sites was allowed. For MS1, a mass tolerance of 10 ppm was allowed, and for MS2, a mass tolerance of 0.6 Da was allowed. A maximum of 3 equal modifications were allowed per peptide, and 4 maximum dynamic modifications were allowed per peptide. For dynamic modifications, oxidation (+15.995 Da) was allowed, and for static modifications, carbamidomethyl (+57.021 Da) was allowed. We used the Percolator node to assign peptide spectral matches (PSMs) and for the decoy database search using a strict FDR of 1%. The Minora Feature Detector node was used to calculate extracted-ion chromatogram (XIC) peak area for quantitation, with a minimum trace length of 5, a minimum number of 2 peaks, and a max *Δ*RT of 0.2 for isotope pattern multiplets.

### Protein quantification

The MSStats package (v. 3.22.1) was used to calculate protein level quantitation from peptide data, as well as to perform differential abundance analysis between fractions and samples (Choi *et al*. 2014). Peptides with only one or two counts across runs were removed, as were proteins with only one peptide. Only unique peptides were used for protein quantitation. Median normalization was performed to normalize extracted ion chromatogram (XIC) peak area across biological replicates and fractions. Protein quantification from peptides was performed using the TOP3 method, and missing values were imputed using an accelerated failure model.

### Differential abundance analysis

The MSStats package (v. 3.22.1) was used to perform differential abundance analysis to identify non-additively expressed proteins. A linear mixed model was used to calculate fold changes and p-values. The mean protein abundance of the hybrid was contrasted against the mean of both parental protein abundance means. Only proteins with measurements in at least two biological replicates per genotype in the soluble fractions were considered. Proteins with p-value ≤ 0.05 and log_2_FC ≥ |0.5| were considered differentially expressed. We used uncorrected p-values as using Benjamini-Hochberg adjusted p-values resulted in the identification of no differentially expressed proteins due to the relatively high variability among the samples. The use of multiple testing correction, although reducing the incidence of Type I errors (false positives), may increase Type II errors (false negatives), as observed in other proteomics studies (Pascovici *et al*. 2016). As a possible remedy, we used a fold-change threshold that may reduce the number of false positives. PCA analysis was performed using the prcomp function in R and drawn using ggbiplot.

### Solubility shift analysis

For this analysis, proteins that were not quantified in all three biological replicates or all fractions were discarded. In base R, a two-way ANOVA, with fraction (insoluble/soluble) and genotype (progenitors/hybrid), and the corresponding interaction term was performed. Proteins with a significant (*P* ≤ 0.05) interaction term displayed a significant shift in solubility between the parents and the hybrid. To quantify the degree to which solubility shifts between the parents and the hybrids, as well as the direction of this shift, a solubility score was calculated. For each biological replicate, the ratio of protein in the soluble fraction to the insoluble fraction was calculated. The median ratio for each progenitor and hybrid was then used for further analysis. Median ratios were used due to the high variability between biological replicates in the insoluble fraction. To get the mid-parent value, the mean was taken of the median ratios for each parent.

The following formula was then used to calculate the overall solubility shift:

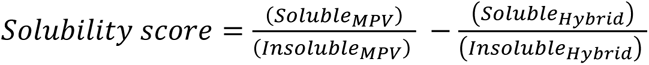

Proteins with p-value ≤ 0.05 and solubility score ≥0.5 were classified as being significantly more soluble in the parents than in the hybrids, and proteins with p-value ≤ 0.05 and solubility score ≤ −0.5 were classified as being significantly more soluble in the hybrids than in the parents.

### Homoeolog-specific protein expression

We assigned peptides to individual homoeologs using the assigned_peptides script from PIVO (https://github.com/marcottelab/pivo) (Drew *et al*. 2020). These peptide matches were then intersected with peptides that were uniquely assigned to an orthogroup. They were filtered to identify peptides that match proteins belonging to either the *A. thaliana* or *A. arenosa* sub-genome for each orthogroup. *A. thaliana* and *A. arenosa* specific peptides were summed separately by orthogroup for each sample. Orthogroups where peptides in the At4 or Aa samples matched to the incorrect parental proteome were discarded. Samples where >75% of peptides matched either the *A. arenosa* or *A. thaliana* proteome were classed as being biased towards that proteome.

### Gene Ontology (GO) analysis

GO analysis was performed using the TopGO package (v. 2.42.0) using the *elim* algorithm (https://bioconductor.org/packages/release/bioc/html/topGO.html). GO annotations for *A. thaliana* from org.At.tair.db were downloaded for enrichment analysis. For the polyploids, orthogroups were annotated by lifting GO annotations from the *A. thaliana* proteins in each orthogroup, using GO annotations downloaded from Ensembl BioMart (Kinsella *et al*. 2011). Orthogroups without an *A.thaliana* member were annotated using InterProScan annotations for the orthogroups assigned in a previous paper (Jiang *et al*. 2021).

### Aggregation propensity and instability predictions

Aggregation propensity was calculated using the TANGO algorithm (Fernandez-Escamilla *et al*. 2004). Instability scores were calculated using the ProtParam tool from Expasy (https://web.expasy.org/protparam).

### RNA-seq analysis

Previously collected RNA-seq data from our lab was used to investigate homoeolog-specific RNA expression in Allo738 and *A. suecica* (NCBI’s Gene Expression Omnibus accession numbers GSE29687 and GSE50715) (Shi *et al*. 2015). Reads were trimmed using trimmomatic (Bolger *et al*. 2014). Reads were then mapped to the Allo738 genome from Jiang et al, 2021, using STAR (Dobin *et al*. 2013) using the following settings--outFilterMismatchNoverLmax 0.04 --outFilterMultimapNmax 20 -- alignIntronMin 25 --alignIntronMax 3000. Reads were then filtered to identify uniquely mapped reads using samtools (using the -q 60 setting) (Barnett *et al*. 2011). For differential expression analysis, reads overlapping each gene were counted using HTseq using the union and reverse stranded settings. EdgeR was used to calculate CPM values for each locus (Robinson *et al*. 2010). Log_2_-fold change (LFC) was calculated between homoeologs within orthogroups. Samples where there was a LFC >2 between homoeologs from either the *A. arenosa* or *A. thaliana* subgenome were classed as “biased.”

## Results

### Proteome in *Arabidopsis* intraspecific hybrids and allotetraploids

We investigated the proteomes of reciprocal hybrids between the *A. thaliana* accessions Col and C24 (Miller *et al*. 2015), a natural allotetraploid *A. suecica* (As), and a resynthesized allotetraploid Allo738, and their progenitors (Wang *et al*. 2006b). *A. thaliana* Col and C24 diverged after the last glacial period (Fig. 1A), about 10,000 years ago (Consortium 2016), while *A. thaliana* (At4) and *A. arenosa* (Aa) diverged around ~6 million years ago and hybridized to form *A. suecica* 16,000-300,000 years ago (Jiang *et al*. 2021; Novikova *et al*. 2017). Both the intraspecific hybrids and allotetraploids display high levels of growth vigor (Fig. 1B), and the level of biomass vigor is higher in the allotetraploids than in intraspecific hybrids, indicating a role of genetic distance in heterosis (Chen 2013; Miller *et al*. 2015).

**Fig 1.**
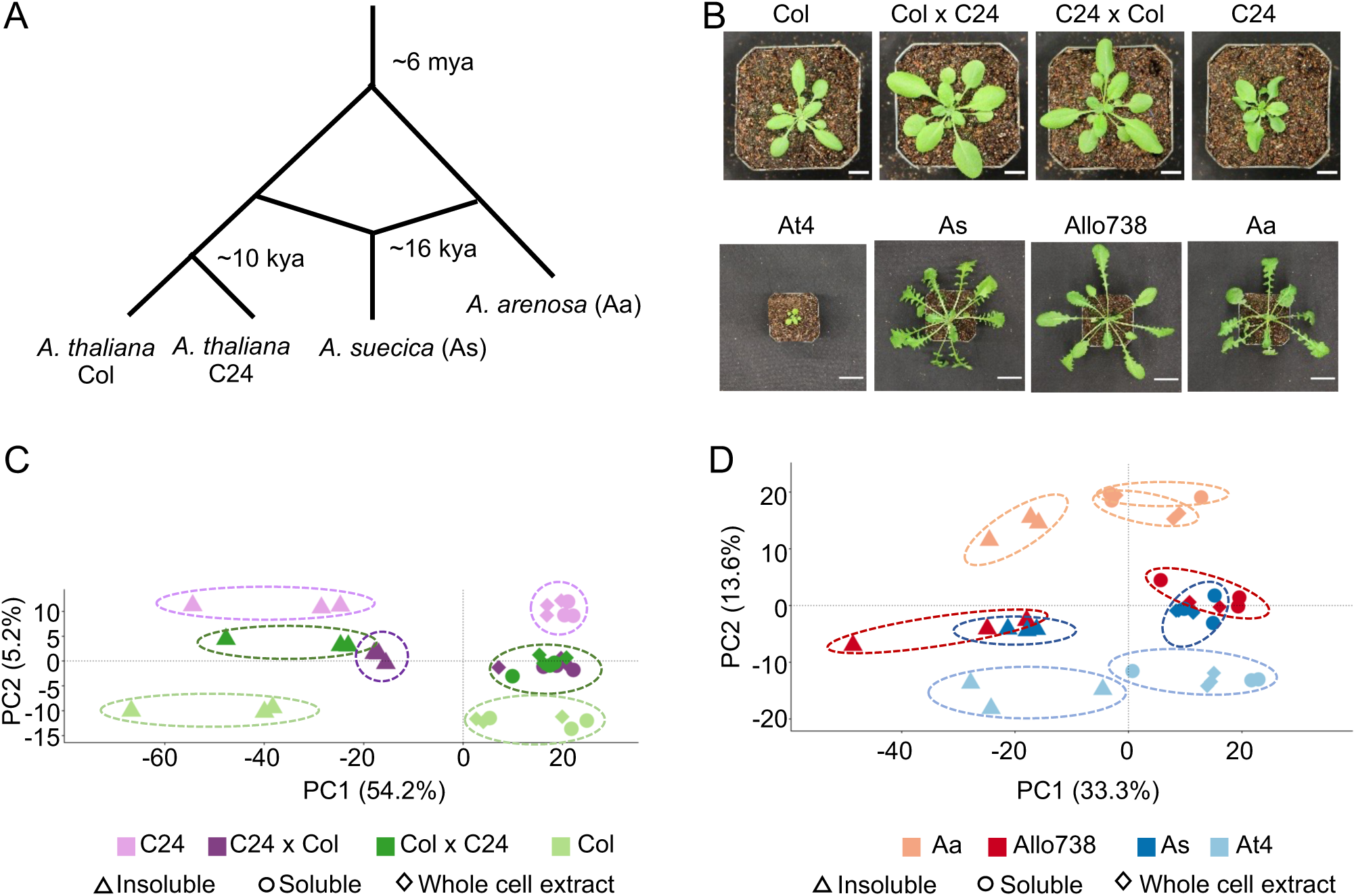
Proteome diversity in Arabidopsis intraspecific hybrids and allotetraploids. (A) Diagram of genetic divergence between *Arabidopsis* species and accessions. Mya: million years ago; kya: thousand years ago. (B) Photographs of *Arabidopsis* intraspecific hybrids and their parents, and of allotetraploids and their (extant) progenitors. Scale bars = 10 mm (intraspecific hybrids) and 30 mm (allotetraploids). (C) PCA plot showing protein abundance identified in the whole cell extract (diamond), soluble fraction (circle), and insoluble (triangle) fractions in the intraspecific hybrids (C24xCol, dark purple and ColxC24, dark green) between C24 (light purple) and Col (light green). There is separation of the samples by genotype along PC2, and separation of the samples by fraction along PC1 with percentage of variation explained (%). (D) PCA plot showing protein abundance in the whole cell extract (diamond), soluble (circle), and insoluble(triangle) fractions in *Arabidopsis* allotetraploids (Allo738, dark red and As, dark blue) and *A. thaliana* (light blue) and *A. arenosa* (light red). As with the intraspecific hybrids, there is separation of the samples by genotype along PC2, and separation of the samples by fraction along PC1.

Native protein extracts were separated into soluble and insoluble fractions using a native and non-denaturing protein extraction method (See Methods), with NP-40 (1%), a non-ionic detergent, at 10,000g centrifugation, and subject to analysis of label-free liquid chromatography mass spectrometry (LC-MS/MS). These proteins were in normal distributions (Supplementary Fig. 1) and highly reproducible among three biological replicates in intraspecific hybrids and their parents (Supplementary Fig. 2) and allotetraploids and their progenitors (Supplementary Fig. 3). We identified a total of 5,144 protein groups (out of 12,769 (Castellana *et al*. 2008)) across all fractions in the intraspecific hybrids, which were filtered down to 2,600 protein groups after removal of non-unique peptides and proteins with few supporting peptides (Supplementary Dataset 1). The recent genome assembly of Allo738 (Jiang *et al*. 2021), comprising the At and Aa sub-genomes, was used to generate a proteome for allotetraploids. To increase protein identifications in the polyploid species, we used an orthogroup-collapsed approach (McWhite *et al*. 2020) for peptide assignment to preserve peptides that were mapped onto both subgenomes in the allotetraploids. The use of this approach had two primary benefits. Firstly, we identified 200 more protein groups and 87,476 more peptide spectrum matches when orthogroups were collapsed than when only the *A. thaliana* proteome was used for peptide assignment. Secondly, it allowed us to evaluate nonadditive expression of proteins in Allo738 and *A. suecica* relative to At4 and Aa. In the allotetraploids, we identified 4,927 protein orthogroups, which were reduced to 2,519 protein orthogroups after removal of lower quality ones (Supplementary Dataset 2). Principal component analysis (PCA) of protein abundance in both *A. thaliana* hybrids (Fig. 1C) and allotetraploids (Fig. 1D) showed clear separation by fractions (PC1) and by genotypes (PC2). In both allotetraploids and intraspecific hybrids, the largest separation was between the two parents Col and C24 for the hybrids (Fig. 1C) and Aa and At4 for the allotetraploids (Fig. 1D), with the hybrids and allotetraploids falling between their respective parents. There was a greater spread along PC2 between allotetraploid progenitors, At4 and Aa (Fig. 1D), than between *A. thaliana* hybrid parents (Col and C24) (Fig. 1C), which could reflect the increased genetic diversity between At4 and Aa compared to Col and C24.

### Proteins are nonadditively expressed in *Arabidopsis* hybrids and allotetraploids

We evaluated protein abundance levels in both allotetraploids and intraspecific hybrids compared to the mid-parent value (MPV) (Supplementary Dataset 3), and differentially expressed proteins between the respective parents (Supplementary Dataset 4). In the intraspecific hybrids, numbers of nonadditively expressed proteins (log_2_FC > |0.5|; p < 0.05) were 109 and 73 in F_1_ (ColxC24, by convention the maternal parent is listed first in a genetic cross) and the reciprocal F_1_ (C24xCol), respectively (Fig. 2A and 2C), and 279 and 228 proteins were nonadditively expressed in As and Allo738, respectively (Fig. 2B and 2D). In the intraspecific hybrids, twice as many proteins that were down-regulated than upregulated, consistent with more down-regulated genes than up-regulated genes in the transcriptome study (Miller *et al*. 2015). However, relatively equal numbers of proteins were upregulated and down-regulated proteins in both allotetraploids, which were inconsistent with microarray data (Wang *et al*. 2006b), but consistent with previous proteomic data (Ng *et al*. 2012). This may suggest a discordance between protein and transcript abundance (Ng *et al*. 2012) and/or different stages of plant materials assayed between two studies.

**Fig 2.**
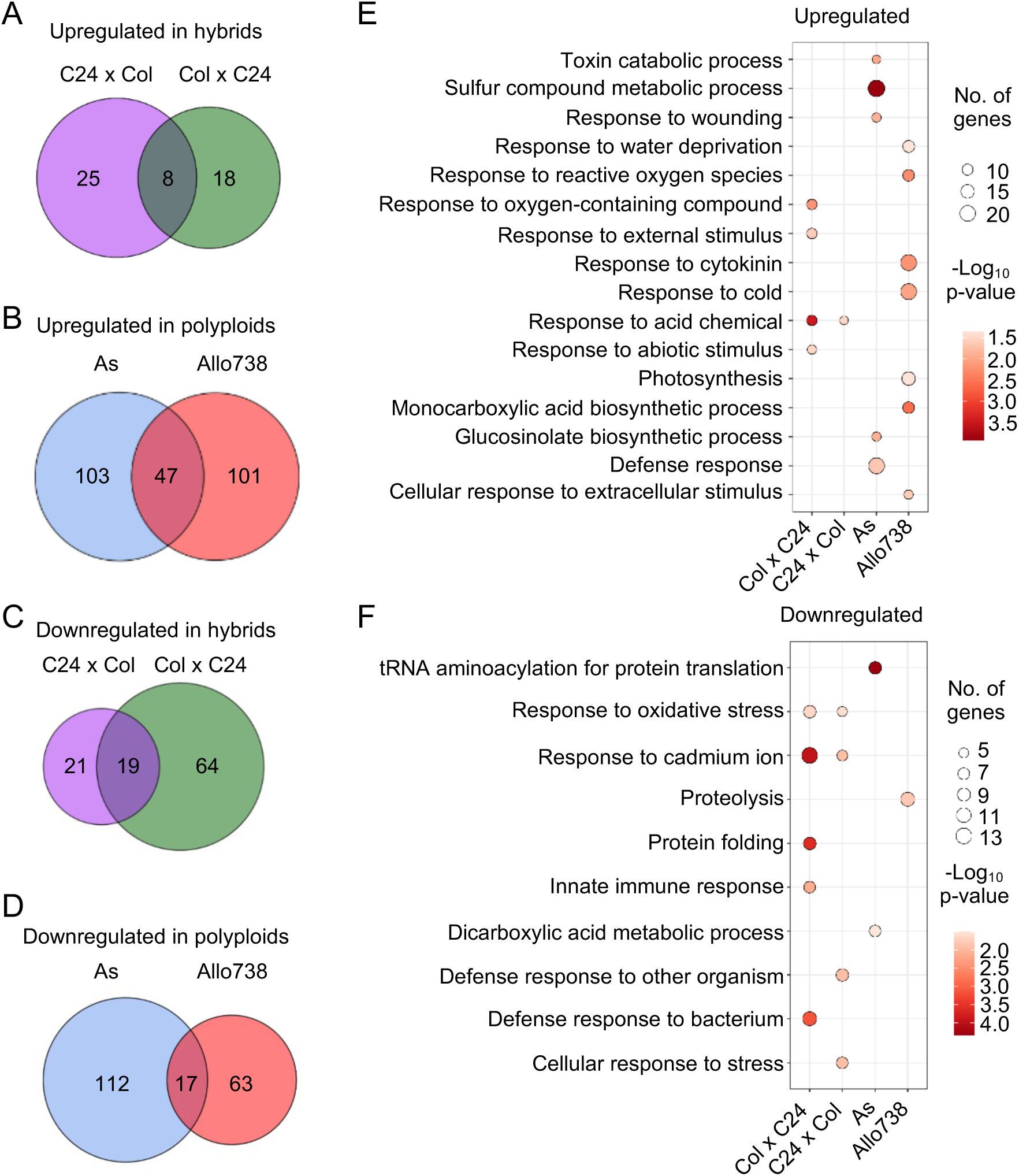
Nonadditive expression of proteins in intraspecific hybrids and allotetraploids. (A) Nonadditively expressed proteins (upregulated) in the hybrids relative to the mid-parent value (MPV). Venn diagrams indicate overlap between Col x C24 and C24 x Col. (B) Nonadditively expressed proteins (upregulated) in the allotetraploids relative to the MPV. Venn diagrams indicate overlap between As and Allo738. (C) Nonadditively expressed proteins (down-regulated) in the hybrids relative to the MPV. Venn diagrams indicate overlap between ColxC24 and C24xCol. (D) Nonadditively expressed proteins (down-regulated) in allotetraploids relative to the MPV. Venn diagrams indicate overlap between Allo738 and As. (E) Gene ontology analysis of biological process enrichment for the upregulated proteins in the hybrids and allotetraploids relative to the MPV. (F) GO analysis of biological process enrichment for the downregulated proteins in the hybrids and allotetraploids relative to the MPV.

The allotetraploids exhibit an increased genetic diversity between their progenitors, as well as an increased level of growth vigor compared to the intraspecific hybrids. This is reflected in the number of nonadditively expressed proteins identified. There was a large degree of overlap in proteins that showed nonadditive expression in the allotetraploids; 98 proteins (43.0%) and 89 proteins (31.9%) were nonadditively expressed in All738 and *A. suecica*, respectively, and were also differentially expressed between *A. thaliana* and *A. arenosa* (Supplementary Fig. 4, A and B). In the F_1_ hybrids (C24xCol and ColxC24), 37 proteins (33.9%) in C24xCol and 27 proteins (37.0%) in ColxC24 were nonadditively expressed and showed differential expression between the parents Col and C24 (Supplementary Fig. 4, C and D). This high-level overlap suggests that protein differences between the parents need to be modified or reconciled in the intraspecific hybrids and allotetraploids, a notion supported by the transcriptome studies (Miller *et al*. 2015; Wang *et al*. 2006b).

### Gene ontology (GO) enrichment of nonadditively expressed proteins

GO analysis identified a number of functional terms as significantly enriched in the nonadditively expressed proteins in both the intraspecific hybrids and allotetraploids (Fig. 2E and 2F). The GO enrichment terms of the nonadditively expressed proteins were much more similar between the two reciprocal intraspecific hybrids than between the allotetraploids. In the intraspecific hybrids, a number of GO enrichment terms were related to stress response, which is consistent with overrepresentation of nonadditively expressed stress-responsive genes in both *Arabidopsis* intraspecific hybrids (Miller *et al*. 2015) and allotetraploids (Wang *et al*. 2006b). Interestingly, the GO enrichment of upregulated proteins was related to the abiotic stress response, such as response to cold (GO:0009409), toxin catabolic process (GO:0009407), and response to acid-containing chemical (GO:1901700) (Fig. 2E), while GO terms of down-regulated proteins were related to the biotic stress response, such as defense response to bacterium (GO:0042742) and defense response to other organism (GO:0098542) (Fig. 2F).

The GO enrichment categories showed little overlap between the nonadditively expressed proteins in As and Allo738, probably because of the large difference between the resynthesized (Allo738) and natural (As) allotetraploids. In Allo738, upregulation of the proteins involved in response to cytokinin (GO:0009735) and cold (GO:0009409) (Fig. 2E) may suggest that cold response is adapted in natural *A. suecica*. Enrichment of GO terms in down-regulated proteins was related to RNA and protein metabolism (Fig. 2F), including tRNA aminoacylation for protein translation (GO:0016070) and proteolysis (GO:0006508). Expression changes of the proteins in protein metabolism agree with previous findings. For example in maize, downregulation of proteins is related to proteasome formation and amino acid biosynthesis (Li *et al*. 2020), and in *Drosophila*, increased inbreeding is associated with an increase in *HSP70* expression (Kristensen *et al*. 2002).

### Analysis of soluble and insoluble proteomes in *Arabidopsis* hybrids and allotetraploids

Theoretical analyses suggest that misfolded proteins can form protein aggregates, leading to proteasomal degradation (Ginn 2010; Ginn 2017) (Fig. 3A). Alternatively, coding sequence variants between alleles in hybrids could lead to a reduction in the rate of self-association during protein folding, leading to a decrease in protein misfolding and aggregation in hybrids (Fig. 3B). To investigate this, we employed a fractionation scheme previously used to investigate protein solubility shifts in *S. cerevisiae* in response to heat shock – in S. cerevisiae proteins enriched in the insoluble fraction were found to form foci after heat shock (O’Connell *et al*. 2014). This method uses a 10,000g centrifugation step to separate the soluble and insoluble fractions from whole cell extract (Fig. 3C).

**Fig 3.**
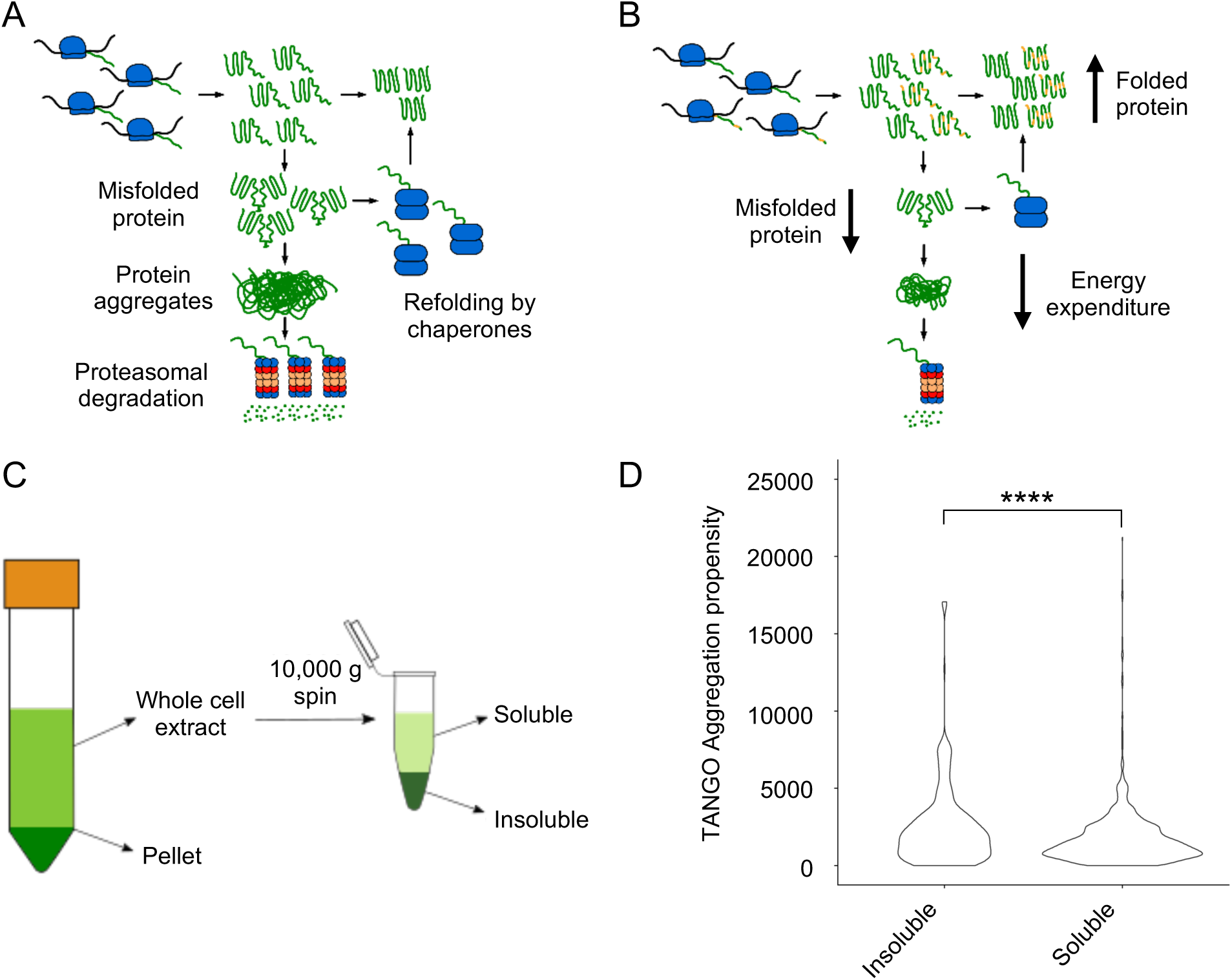
Soluble and soluble fractions of proteomes in hybrids and allotetraploids. (A) A model of protein aggregation in inbred lines. Homo-oligomerization occurs while proteins fold after synthesis, and these either form larger aggregates, which are degraded by the proteasome or refolded by chaperone proteins. (B) A model of protein aggregation in hybrids that could explain their increased metabolic efficiency observed in hybrids. Changes in protein coding sequence between the two alleles of a gene (represented in yellow) could prevent protein aggregation by disrupting homo-oligomerization, reducing the amount of aggregate and thus the proteins that must be degraded or refolded. (C) Fractionation scheme from low (1000 g, left) to high (10,000 g, right) speed to isolate the soluble and insoluble proteomes, respectively. (D) Distribution of TANGO aggregation propensity scores between the soluble and insoluble fractions. Four asterisks indicate the statistical significance level of *P*<0.0001 (Wilcoxon Rank Sum test).

This method was previously successful in separating insoluble and soluble proteins in *Arabidopsis* on the basis of aggregation propensity as calculated using the TANGO algorithm (Fernandez-Escamilla *et al*. 2004). We therefore evaluated whether there was a significant difference in the TANGO scores of proteins enriched in the soluble and insoluble fractions of the proteome in our samples. The insoluble fraction had proteins with a significantly higher mean TANGO score than the proteins in the soluble fraction (*P* = 1.36 x 10^-7^, Wilcoxon Rank Sum test), indicating that it is enriched in aggregating proteins (Fig. 3D).

GO analysis found that proteins more abundant in the soluble fraction were represented many cellular components and most cell regions, whereas proteins more abundant in the insoluble fraction only showed enrichment in a few cellular components primarily membrane-bound organelles such as the chloroplast envelope (GO:0009941) and the thylakoid membrane (GO:0009535) (Supplementary Fig. 5).

Cytosolic proteins, such as ribosomal proteins, were generally enriched in the soluble fraction instead of the insoluble fraction. As with the insoluble fraction, there was an enrichment of chloroplast localized proteins; however, unlike the insoluble fraction, there was enrichment of proteins from the thylakoid lumen and the stroma in addition to the thylakoid membrane. This included both subunits of RuBisCO, which were significantly enriched in the soluble fraction of all samples. This argues that the abundance of thylakoid proteins in the chloroplast in the insoluble fraction is not due to intact chloroplasts accumulating in the insoluble fraction, but rather a reflection of the solubility of these proteins. This finding may suggest a role for protein solubility in maintaining high photosynthetic activities, as they contribute to heterosis in *Arabidopsis* (Ni *et al*. 2009) and maize (Ko *et al*. 2016; Li *et al*. 2020).

### Changes in protein solubility between hybrids and their parents

To test a potential role of protein solubility changes in hybrid vigor, we examined whether there was a general shift in protein solubility of proteins between hybrids and their parents. Using ANOVA (*P* < 0.05), we calculated pairwise ratios of the protein abundance between soluble and insoluble fractions. When the median ratio was greater than 0.5 between the MPV ratio and hybrid ratio in addition to a *P*-value of less than 0.05, the proteins were considered having a solubility shift between the hybrids and the parents (Fig. 4A) (Supplementary Dataset 5).

**Fig 4.**
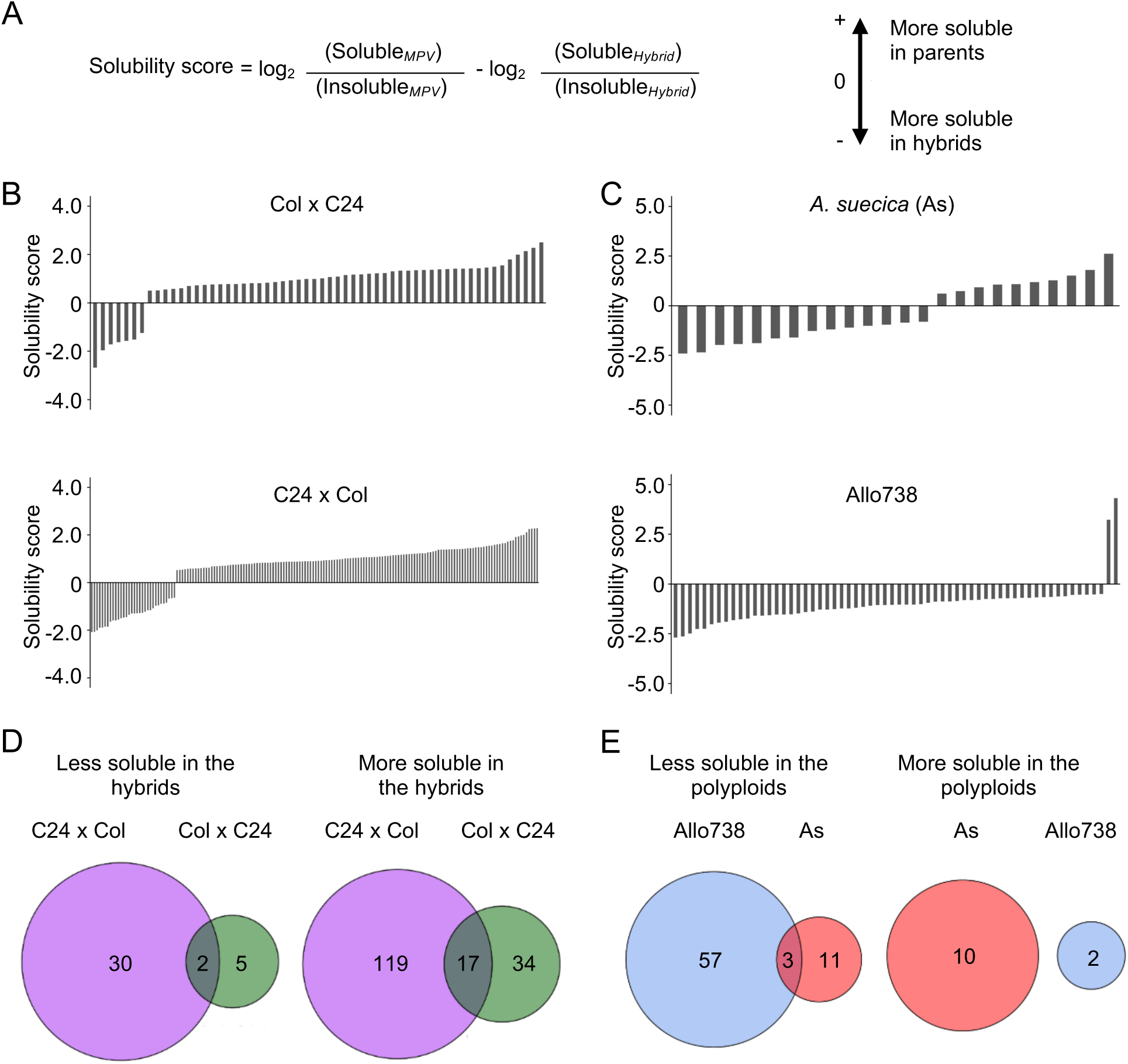
Analysis of protein solubility in intraspecific hybrids and allotetraploids. (A) The solubility score calculated to identify whether the proteins increased or decreased in solubility between a hybrid or polyploid and the parents. (B) Solubility scores of proteins in Col x C24 and C24 x Col. (C) Solubility scores of proteins in *A. suecica* (As) and Allo 738. (D) Venn diagrams displaying overlap in proteins that change solubility in hybrids between Col x C24 and C24 x Col (E) Venn diagrams displaying overlap in proteins that change solubility in allotetraploids Allo738 and As.

In the intraspecific hybrids relative to the parents, there were more proteins in the soluble than in the insoluble fractions (Fig. 4B and 4D). Among those soluble proteins that were localized in the chloroplast stroma, 14 out of 34 proteins were more soluble in both reciprocal hybrids than in their parents. Col x C24 hybrids were shown to have more chloroplasts per cell and an increased photosynthetic capacity (Fujimoto *et al*. 2012), which could contribute to the increase in protein solubility observed for these proteins. In both hybrids, fewer proteins showed lower solubility than in their parents. Only two proteins were less soluble in both hybrids: *RBP31*, a chloroplast ribonucleoprotein, and *OEP16*, a chloroplast outer envelope pore protein.

Unexpectedly, fewer proteins displayed a solubility shift in the allotetraploids than in the intraspecific hybrids (Fig. 4B and 4C). Ten and two proteins were more soluble in Allo738 and As, respectively (Fig. 4E), compared to their progenitors, including a heat shock factor binding protein that is involved in acquired thermotolerance (Hsu *et al*. 2010). Three protein orthogroups showed a reduced solubility relative to both progenitors: an outer envelope membrane protein, a hydroxymethylglutaryl-CoA synthase involved in glucosinolate biosynthesis and an orthogroup containing kinesin-like protein involved in cell division. In addition, there were more proteins that showed a decrease in protein solubility in the allotetraploids relative to their progenitors. These data may suggest protein solubility may not be directly related to genetic distance. Alternatively, protein solubility may change during different stages of development, as these allotetraploids grow slower and flower later than the diploids (Wang *et al*. 2006a).

### Expression of homoeolog-specific proteins in allotetraploids

The recent genome assembly for Allo738 (Jiang *et al*. 2021) allowed us to use the Allo738 proteome for peptide assignments. This improved reference proteome, along with the increased divergence between At4 and Aa helped us identify peptides that were unique to individual homoeologs in the allotetraploids (Supplementary Dataset 6). This would allow us to test if allelic-specific expression of proteins in hybrids and polyploids contributes to the metabolic efficiency in hybrids (Goff 2011).

Allele-specific peptides were used to calculate protein abundance in allotetraploids, and the abundance of proteins from the Aa and At sub-genomes were compared within each orthogroup. We found that similar numbers of proteins that displayed a bias towards either the *A. thaliana* or *A. arenosa* subgenome in Allo738 and natural *A. suecica* (Fig. 5, A and C); nearly 50% of these proteins in At-biased (Fig. 5A) or Aa-biased (Fig. 5C) group were shared between Allo738 and natural *A. suecica*. This finding is consistent with transcriptome data that no obvious expression dominance was found among multiple natural *A. suecica* accessions (Burns *et al*. 2021). Although expression dominance of specific homoeologs can occur in the allotetraploids *Arabidopsis* (Wang *et al*. 2006a; Wang *et al*. 2006b). cotton (Adams *et al*. 2003; Zhang *et al*. 2015), and *Tragopogon* (Tate *et al*. 2006), our data support the notion of genomic and expression stability accompanied by epigenetic changes in many genetically stable allopolyploids like *Arabidopsis* (Jiang *et al*. 2021) and *Gossypium* (cotton) (Chen *et al*. 2020).

**Fig 5.**
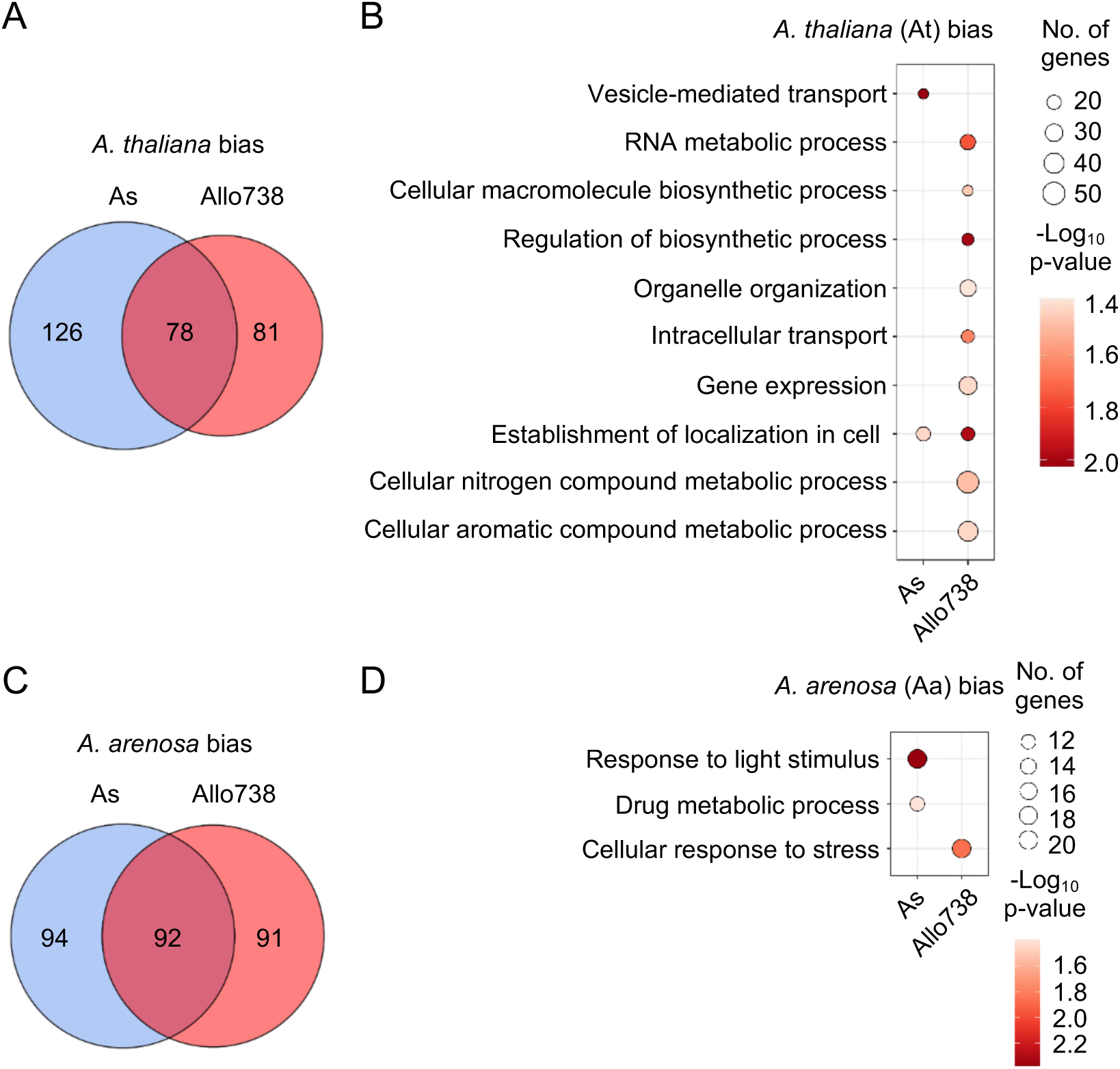
Homoeolog-specific expression of proteins in allotetraploids. (A) Venn diagram indicating the overlap between proteins that display biased expression towards the *A. thaliana* homoeologs in Allo738 and natural *A. suecica* (As). (B) Enrichment of GO biological process for proteins that display *A. thaliana* homoeolog*-*biased expression in Allo738 and As. (C) Venn diagram indicating the overlap between proteins that display biased expression of the *A. arenosa* homoeologs in Allo738 and As. (D) Enrichment of GO biological process for proteins that display *A. arenosa* homoeolog*-*biased expression in Allo738 and As.

Among orthogroup proteins that display biased-homoeolog expression, *A. thaliana*-biased proteins showed more GO enrichment groups than *A. arenosa*-biased proteins (Fig. 5, B and D). At-biased proteins had more GO enrichment terms in Allo738 than natural *A. suecica* (As), many of which belonged to biosynthetic and metabolic processes, including RNA metabolic process (GO:0016070), organelle organization (GO:0006996), regulation of biosynthetic process (GO:0009889), gene expression (GO:0010467), and cellular nitrogen and aromatic compound metabolic processes. Many of these proteins are localized in chloroplasts; this may reflect inheritance of chloroplasts from the maternal *A. thaliana* ancestor of Allo738 (Wang *et al*. 2006b) and *A. suecica* (Sall *et al*. 2003). Alternatively, proteins, like transcripts, of *A. thaliana* origin, may be subject to biased expression (Wang *et al*. 2006b). The *A. arenosa*-biased proteins had fewer GO enrichment terms, including response to light stimulus in both Allo738 and As, and the cellular response to stress in Allo738.

These homoeolog-biased proteins accounted for 20% of nonadditively expressed proteins in *A. suecica* and 17% in Allo738 (Supplementary Dataset 6), indicating a role of homoeolog-biased expression in the nonadditive protein accumulation in allotetraploids. Interestingly, about twice as many nonadditively expressed proteins that displayed homoeolog-expression bias were expressed above MPV than below MPV in As, despite the fact that the percentage of nonadditively expressed proteins displaying homoeolog-expression bias was similar in both allotetraploids. This contrasted with Allo738, in which equal numbers of nonadditively expressed proteins were expressed both above and below MPV. This may reflect changes in protein abundance (or silencing) between neo-allopolyploid Allo738 and natural *A. suecica*.

To determine whether homoeolog-expression bias contributes to changes in protein solubility in hybrids, we estimated the mean TANGO score and instability score (Supplementary Fig. 3) and compared them for both homoeologous proteins that displayed biased expression. If there was a trend towards expressing the more stable homoeologs, we would expect to see an increase in the average solubility of At or Aa homoeologs. However, no obvious difference was observed in the aggregation propensity of the proteins displaying biased expression in either Allo738 or *A. suecica* (Supplementary Fig. 6) and transcriptome level (Supplementary Fig. 7). Homoeolog-expression bias may not alter protein solubility in *Arabidopsis* allotetraploids.

## Discussion

### Nonadditive accumulation of stress-response proteins in intraspecific hybrids and allotetraploids

Our investigation of the proteome uncovered the role of non-additively expressed proteins in stress responses in *Arabidopsis* intraspecific hybrids and allotetraploids. Down-regulation of abiotic and biotic stress-responsive genes in normal conditions can save the energy to promote growth vigor (Miller *et al*. 2015). Results from the proteomic analysis largely support the findings of nonadditively expressed transcripts in *Arabidopsis* intraspecific hybrids (Miller *et al*. 2015) and allotetraploids (Wang *et al*. 2006b). For example, the biotic stress-responsive proteins that are downregulated include two proteins in the *PATHOGENESIS-RELATED GENES* family, *PR2* and *PR5*. Members of this gene family are also downregulated at mRNA levels in Col/C24 hybrids (Miller *et al*. 2015). Several genes encoding glutathione-S-transferase (GST), *GSTF6* and *GSFT2*, are downregulated in both reciprocal hybrids, and *GSTF7* and *GSTF8* are downregulated in one F_1_ (Col x C24). *GST* genes are involved in response to bacterial or fungal infections by removing toxins associated with pathogen infection as well as in mediating a systemic immune response (Gullner *et al*. 2018).

Moreover, a reduction in oxidative stress can potentially downregulate protein metabolic machinery. In maize hybrids, catalase protein abundance is greater than mid-parent value at ZT21, leading to an overnight reduction in H_2_O_2_ abundance (Li *et al*. 2020). In the allotetraploids, the orthogroup containing catalase gene orthologs (OG0000577) is significantly upregulated in Allo738 relative to the mid-parent value (*P* = 0.013) and slightly in *A. suecica* (*P* = 0.053). This increased expression of catalase in the allotetraploids may contribute to low levels of protein damage due to oxidative stress, thus leading to a reduction in the requirement of protein biosynthesis machinery in hybrids and polyploids.

Among the proteins upregulated relative to the MPV in both Allo738 and As, many are related to photosynthesis, consistent with upregulation of these genes in resynthesized allotetraploids (Ni *et al*. 2009; Wang *et al*. 2006b). For example, AMY3, an alpha amylase protein involved in starch degradation, is upregulated in both allotetraploids, and its transcripts are also upregulated in resynthesized allotetraploids (Ni *et al*. 2009; Wang *et al*. 2006b). In addition, PORC, a protochlorophyllide oxidoreductase that is involved in the biosynthesis of chlorophyll, is upregulated in both allotetraploids. Other POR loci, such as *PORA* and *PORB* are also found to be consistently upregulated in allotetraploids (Ni *et al*. 2009; Wang *et al*. 2006b).

Upregulated proteins of *A. thaliana* homoeologs in the allotretraploids include β-glucosidases and jacalin-related lectin *JAL35*, which is involved in glucosinolate biosynthesis and ER body formation (Nagano *et al*. 2008). ER bodies are responsible for the formation of isothiocyanates, which are toxic to many herbivores (Wittstock *et al*. 2003). Upregulation of these proteins in the allotetraploids may contribute to glucosinolate turnover pathway. Furthermore, the *A. thaliana* homoeolog of the GRP7 protein orthogroup is upregulated in both Allo738 and *A. suecica*, consistent with microarray results (Wang *et al*. 2006b). GRP7 is an RNA binding protein that is involved in regulating circadian oscillation (Heintzen *et al*. 1997), as well as both biotic and abiotic stress-responsive genes (Meyer *et al*. 2017). Upregulation of this protein could mediate expression of stress-responsive genes by altering the circadian clock in the intraspecific hybrids and allotetraploids (Miller *et al*. 2015; Ni *et al*. 2009).

### Cytokinin responsive proteins were non-additively expressed in the allotetraploids

Phytohormones, including the ethylene, salicylic acid, and auxin, have been shown to play roles in mediating growth vigor in hybrids (Groszmann *et al*. 2015; Saeki *et al*. 2016; Shen *et al*. 2012; Song *et al*. 2018; Wang *et al*. 2006b). In this study, we found upregulation of genes involved in response to cytokinin in the allotetraploids. Upregulation of the proteasome subunit RPN12a in Allo738 may be involved in promoting the degradation of inhibitors to the cytokinin response. For example, the *RPN12a* mutant shows slow leaf formation, reduced root elongation, and altered growth in response to exogenous cytokinins (Smalle *et al*. 2002). Cytokinins are generally involved in promoting cell division and plant growth – mutants that overexpress cytokinin biosynthesis genes are associated with increased shoot growth (Kieber and Schaller 2014). This suggests that cytokinin may play a role in mediating hybrid vigor in the allotetraploids.

### Changes in protein solubility in *Arabidopsis* intraspecific hybrids and allotetraploids

Theoretical studies of protein folding in yeast hybrids suggest hybrids have lower levels of protein aggregation, and thus more soluble proteins (Ginn 2010; Ginn 2017). Consistent with this, downregulation of genes involved in protein metabolism is observed in intraspecific hybrids in *Drosophila* (Kristensen *et al*. 2002). Here we found a shift of protein solubility in the intraspecific hybrids relative to the mid-parent value, but not in the allotetraploids. It is possible that factors other than genetic distance affect the observed changes in solubility. Alternatively, a computational estimate of a protein’s solubility in yeast hybrids may not reflect its stability *in vivo*. We also note that although the method of separating soluble and insoluble proteins has been successfully used in yeast studies (O’Connell *et al*. 2014), it should be refined for working with plant cells that have rigid cell walls and more debris than the yeast cells. Moreover, appropriate statistical methods and additional validation are needed to properly interpret these data.

Our results also confirm the enrichment of several components of the TIC-TOC complex (translocon on the inner chloroplast membrane - translocon on the inner chloroplast membrane), as well as many members of the photosystem II reaction center in the insoluble fraction in all samples. These proteins are largely located in the chloroplast’s membranes and less soluble than cytosolic proteins. TIC214, a component of the TIC-TOC complex having amyloidogenic properties due to the QN-rich region of the protein’s C-terminus (Antonets and Nizhnikov 2017), is significantly enriched in the insoluble fraction of all samples where it was detected. Many of these proteins are highly abundant with about 80% of protein molecules in a mesophyll cell localized to the chloroplast (Heinemann *et al*. 2021).

In analysis of 400 and 350 proteins with homoeolog-specific expression in *A. suecica* and Allo738, respectively, we did not observe any significant differences in aggregation propensity of the homoeologs. This result is consistent with overall balanced expression among subgenomes (Jiang *et al*. 2021), despite expression bias can occur to rRNA genes and other protein-coding genes due to epigenetic changes (Chen *et al*. 1998; Lee and Chen 2001).

What might cause the increase in metabolic efficiency and downregulation of genes related to protein biosynthesis? One possibility is novel functionality of protein complexes emerging from protein-protein interactions between different protein alleles in the hybrids (Herbst *et al*. 2017) or in the allotetraploids. In *S. cerevisiae* x *S. uvarum* hybrids, there is an overrepresentation of proteins involved in protein metabolism that displayed protein-protein interactions between diverged alleles (Berger and Landry 2021; Dandage *et al*. 2021). A number of complexes involved in protein metabolism, including the prefoldin complex and proteasome, consist of members from both parental copies. This is reminiscent of the abundance of metabolites and proteins in hybrid maize, where most amino acids show abundance peaks during the day and falling at night (Li *et al*. 2020). In addition, investigation of the circadian control of protein synthesis in both *Arabidopsis* and dinoflagellates has found that ribosome loading and translation primarily occurs overnight (Cornelius *et al*. 1985; Missra *et al*. 2015). This may lead to the decreased expression in amino acid biosynthesis and tRNA synthesis as observed in this study and in maize hybrids (Li *et al*. 2020). Whether novel protein-protein interactions have altered function that could impact proteostasis in hybrids and hybrid vigor remains to be investigated.

## Data availability

All raw and interpreted mass spectrometry data were deposited to the ProteomeXchange https://massive.ucsd.edu with the MassIVE repository number MSV000089682 and ProteomeXChange number PXD034635.

## Acknowledgements

We thank Dr. Alan Lloyd at The University of Texas at Austin for supervision in the latter part of this project and Texas Advanced Computing Center for providing computing support for data analysis.

## Author contributions

V.J. and Z.J.C. conceived the research, analyzed the data, and wrote the paper. V.J., D.X., O.P., and D.B. performed the experiments. E.M.M. provided supervision, revision, and technical and intellectual support.

## Supporting Information

### Supplemental Figures

**Supplementary Fig. 1. Log_2_(normalized precursor abundance) values of the peptides among all samples analyzed.** Precursor ion abundance is calculated based on the chromatogram peak area of the precursor ion. (A) Log_2_(normalized precursor abundance) of all peptides identified in the intraspecific hybrid samples (B) Log_2_(normalized precursor abundance) of all peptides identified in the interspecific polyploid samples.

**Supplementary Fig. 2. Reproducibility of protein abundance in intraspecific hybrids Samples.** Pearson correlation of protein abundances for all intraspecific hybrid samples. There is a high degree of reproducibility between biological replicates, with the insoluble fractions showing lower levels of correlation with soluble and whole cell extract fractions.

**Supplementary Fig. 3. Reproducibility of protein abundance in allotetraploids**. Pearson correlation of protein abundances for all allotetraploid samples. There is a high degree of reproducibility between biological replicates, with the insoluble fractions showing lower levels of correlation with soluble and whole cell extract fractions.

**Supplementary Fig. 4. Overlap of the proteins that were differentially expressed between the parents with the non-additively expressed proteins.** The proteins differentially expressed between the progenitors of each hybrid or allotetraploid were overlapped with the non-additively expressed proteins in (A) All738, (B) *A. suecica*, (C) F_1_ (Col x C24), and (D) F_1_ (C24 x Col).

**Supplementary Fig. 5. Enrichment of soluble fractions in cytosolic proteins and of insoluble fractions of proteins localized to membranes.** GO cellular component enrichment of proteins significantly enriched in the insoluble and soluble fractions in both experiments. The soluble fraction shows strong enrichment across several cellular components including the cytosol, whereas the insoluble fraction only shows enrichment for chloroplast proteins, particularly those in the thylakoid membrane.

**Supplementary Fig. 6. Evaluation of homoeolog-specific expression found no obvious homoeolog-specific expression of more soluble proteins in *Arabidopsis* allotetraploids.** The distribution of TANGO scores of the *A. thaliana* and *A. arenosa* homoeologs of all proteins that display homoeolog biased expression in (A) Allo738 and (B) *A. suecica* and the distribution of instability scores of the *A. thaliana* and *A. arenosa* homoeologs of all proteins that display homoeolog biased expression in (C) Allo738 and (D) *A. suecica*. For each graph, grey boxes divide graph into proteins that displayed either *A. thaliana* or *A. arenosa* subgenome bias, and the two distributions within these panels indicate the distribution of TANGO or instability scores for the *A. arenosa* and *A. thaliana* homoeologs of the proteins that display this biased expression.

**Supplementary Fig. 7. Evaluation of homoeolog-specific expression found no obvious homoeolog-specific expression of more soluble proteins in *Arabidopsis* allotetraploids.** The distribution of TANGO scores of the *A. thaliana* and *A. arenosa* homoeologs of all proteins that display homoeolog biased expression in (A) Allo738 and (B) *A. suecica* and the distribution of instability scores of the *A. thaliana* and *A. arenosa* homoeologs of all proteins that display homoeolog biased expression in (C) Allo738 and (D) *A. suecica*. For each graph, grey boxes divide graph into proteins that displayed either *A. thaliana* or *A. arenosa* subgenome bias, and the two distributions within these panels indicate the distribution of TANGO or instability scores for the *A. arenosa* and *A. thaliana* homoeologs of the proteins that display this biased expression.

### Supplemental Datasets

**Supplementary Dataset 1. Protein-level quantification in intraspecific hybrids and their parents**

**Supplementary Dataset 2. Protein-level quantification in allotetraploids and their progenitors**

**Supplementary Dataset 3. Nonadditively expressed proteins in allotetraploids and intraspecific hybrids**

**Supplementary Dataset 4. Differentially expressed proteins between progenitors**

**Supplementary Dataset 5. Solubility shift of proteins between hybrids and their parents**

**Supplementary Dataset 6. Homoeolog-specific expression of proteins in allotetraploids**

